# Ecology Shapes Microbial Immune Strategy: Temperature and Oxygen as Determinants of the Incidence of CRISPR Adaptive Immunity

**DOI:** 10.1101/326330

**Authors:** Jake L. Weissman, Rohan M. R. Laljani, William F. Fagan, Philip L. F. Johnson

## Abstract

Bacteria and archaea are locked in a near-constant battle with their viral pathogens. Despite previous mechanistic characterization of numerous prokaryotic defense strategies, the underlying ecological drivers of different strategies remain largely unknown and predicting which species will take which strategies remains a challenge. Here, we focus on the CRISPR immune strategy and develop a phylogenetically-corrected machine learning approach to build a predictive model of CRISPR incidence using data on over 100 traits across over 2600 species. We discover a strong but hitherto-unknown negative interaction between CRISPR and aerobicity, which we hypothesize may result from interference between CRISPR associated proteins and non-homologous end-joining DNA repair due to oxidative stress. Our predictive model also quantitatively confirms previous observations of an association between CRISPR and temperature. Finally, we contrast the environmental associations of different CRISPR system types (I, II, III) and restriction modification systems, all of which act as intracellular immune systems.

## Introduction

In the world of prokaryotes, infection by viruses poses a constant threat to continued existence (e.g., [1]). In order to evade viral predation, bacteria and archaea employ a range of defense mechanisms that interfere with one or more stages of the viral life-cycle. Modifications to the host’s cell surface can prevent viral entry in the first place. Alternatively, if a virus is able to enter the host cell, then intracellular immune systems, such as the clustered regularly inter-spaced short palindromic repeat (CRISPR) adaptive immune system or restriction-modification (RM) innate immune systems, may degrade viral genetic material and thus prevent replication [2, 3, 4, 5, 6, 7]. Despite our increasingly in-depth understanding of the mechanisms behind each of these defenses, we lack a comprehensive understanding of the factors that cause selection to favor one defense strategy over another.

Here we focus on the CRISPR adaptive immune system, which is a particularly interesting case study due to its uneven distribution across prokaryotic taxa and environments. Previous analyses have shown that bacterial thermophiles and archaea (both mesophilic and thermophilic) frequently have CRISPR systems (~90%), whereas less than half of mesophilic bacteria have CRISPR (~40%; [8, 9, 10, 11, 12]). Environmental samples have revealed that many uncultured bacterial lineages have few or no representatives with CRISPR systems, and that the apparent lack of CRISPR in these lineages may be linked to an obligately symbiotic lifestyle and/or a highly reduced genome [13]. Nevertheless, no systematic exploration of the ecological conditions that favor the evolution and maintenance of CRISPR immunity has been made. Additionally, though these previous results appear broadly true [14], no explicit accounting has been made for the potentially confounding effects of phylogeny in linking CRISPR incidence to particular traits.

What mechanisms might shape the distribution of CRISPR systems across microbes? Some researchers have emphasized the role of the local viral community, suggesting that when viral diversity and abundance is high CRISPR will fail, and thus be selected against [11, 12, 15]. Others have focused on the tradeoff between constitutively expressed defenses like membrane modification and inducible defenses such as CRISPR [15]. Yet others have noted that hot, and possibly other extreme environments can constrain membrane evolution, necessitating the evolution of intracellular defenses like CRISPR or RM systems [16, 17, 18]. Many have observed that since CRISPR prevents horizontal gene transfer, it may be selected against when such transfers are beneficial (e.g. [19, 20]). More recently it has been shown that at least one CRISPR-associated (Cas) protein can suppress non-homologous end-joining (NHEJ) DNA repair, which may lead to selection against having CRISPR in some taxa [21]. In order to determine the relative importances of these different mechanisms, we must first identify the habitats and microbial lifestyles associated with CRISPR immunity.

Here we aim to expand on previous analyses of CRISPR incidence in three ways: (1) by drastically expanding the number of environmental and lifestyle traits considered as predictors using the combination of a large prokaryotic trait database and machine learning approaches, (2) by incorporating appropriate statistical corrections for non-independence among taxa due to shared evolutionary history, and (3) by simultaneously looking for patterns in RM systems, which will help us untangle the difference between environments that specifically favor CRISPR adaptive immunity versus DNA-degrading intracellular immune systems in general (RM and CRISPR).

## Results

Below, we associate specific microbial immune strategies with a diverse list of microbial traits. The traits span a range of scales including aspects of habitat (e.g. “aquatic”), morphology (e.g., “coccus”), and physiology (e.g., “heterotroph”) [44]. While this variety of scales poses a modeling challenge to traditional approaches including linear regression, machine learning algorithms provide an elegant means of integrating such multi-scale traits in a statistically rigorous predictive framework. In particular, we apply algorithms that excel at identifying both linear and non-linear combinations of traits with high predictive ability.

### Visualizing CRISPR Incidence in Trait Space

We visualized CRISPR incidence in microbial trait space using two unsupervised algorithms to collapse high-dimensional data (174 binary traits assessed in 2679 species; see Methods) into fewer dimensions. Both methods revealed clear differences between the placement of CRISPR-encoding and CRISPR-lacking organisms in trait space, despite the fact that no explicit information about CRISPR was included.

First, principal components analysis (PCA) of the trait data reveals several well accepted patterns of microbial lifestyle choice and CRISPR incidence. The first principal component (17% variance explained) corresponds broadly to an axis running from host-associated to free-living microbes (Table 1), as observed by others [23, 24]. CRISPR-encoding and CRISPR-lacking microbes are not differentiated along this axis (S1 Fig). We see CRISPR-encoding and CRISPR-lacking organisms beginning to separate along the second (10% variance explained) and third (7% variance explained) principal components (Fig 1). The second component roughly represents a split between extremophilic, energy-stressed species and mesophilic, plant-associated species (Table 1). Optimal growth temperature appears to be an important predictor of CRISPR incidence, as previously noted by others [11, 12]. The third component is not as easy to interpret, but appears to indicate a spectrum from group living microbes (e.g. biofilms) to microbes that tend to live as lone, motile cells (Table 1). That CRISPR is possibly favored in group-living microbes is not entirely surprising, considering the increased risk of viral outbreak at high population density, and that some species up-regulate CRISPR during biofilm formation [25].

**Table 1:** Top 10 variable loadings on the first three principal components of the microbial traits dataset. These three components explain 17%, 10%, and 7% of the total variance, respectively.

| PC1 | Weight | PC2 | Weight | PC3 | Weight |
| --- | --- | --- | --- | --- | --- |
| ecosystemcategory_human | -0.16 | temperaturerange_mesophilic | 0.19 | growth_in_groups | -0.24 |
| specificecosystem_sediment | 0.16 | temperaturerange_thermophilic | -0.19 | gram_stain_positive | -0.24 |
| ecosystem_environmental | 0.16 | oxygenreq_strictanaero | -0.19 | cellarrangement_singles | 0.21 |
| knownhabitats_host | -0.15 | temperaturerange_hyperthermophilic | -0.18 | cellarrangement_filaments | -0.20 |
| ecosystemsubtype_intertidalzone | 0.15 | knownhabitats_hotspring | -0.17 | sporulation | -0.20 |
| ecosystem_hostassociated | -0.15 | exosystemtype_rhizoplane | 0.17 | energysource_chemoorganotroph | -0.19 |
| habitat_hostassociated | -0.15 | habitat_specialized | -0.16 | cellarrangement_clusters | -0.18 |
| habitat_freeliving | 0.15 | metabolism_methanogen | -0.16 | shape_tailed | -0.18 |
| ecosystemtype_digestivesystem | -0.14 | ecosystemcategory_plants | 0.15 | habitat_terrestrial | -0.18 |
| specificecosystem_fecal | 0.14 | ecosystemtype_thermalsprings | -0.15 | motility | 0.17 |

**Figure 1:**
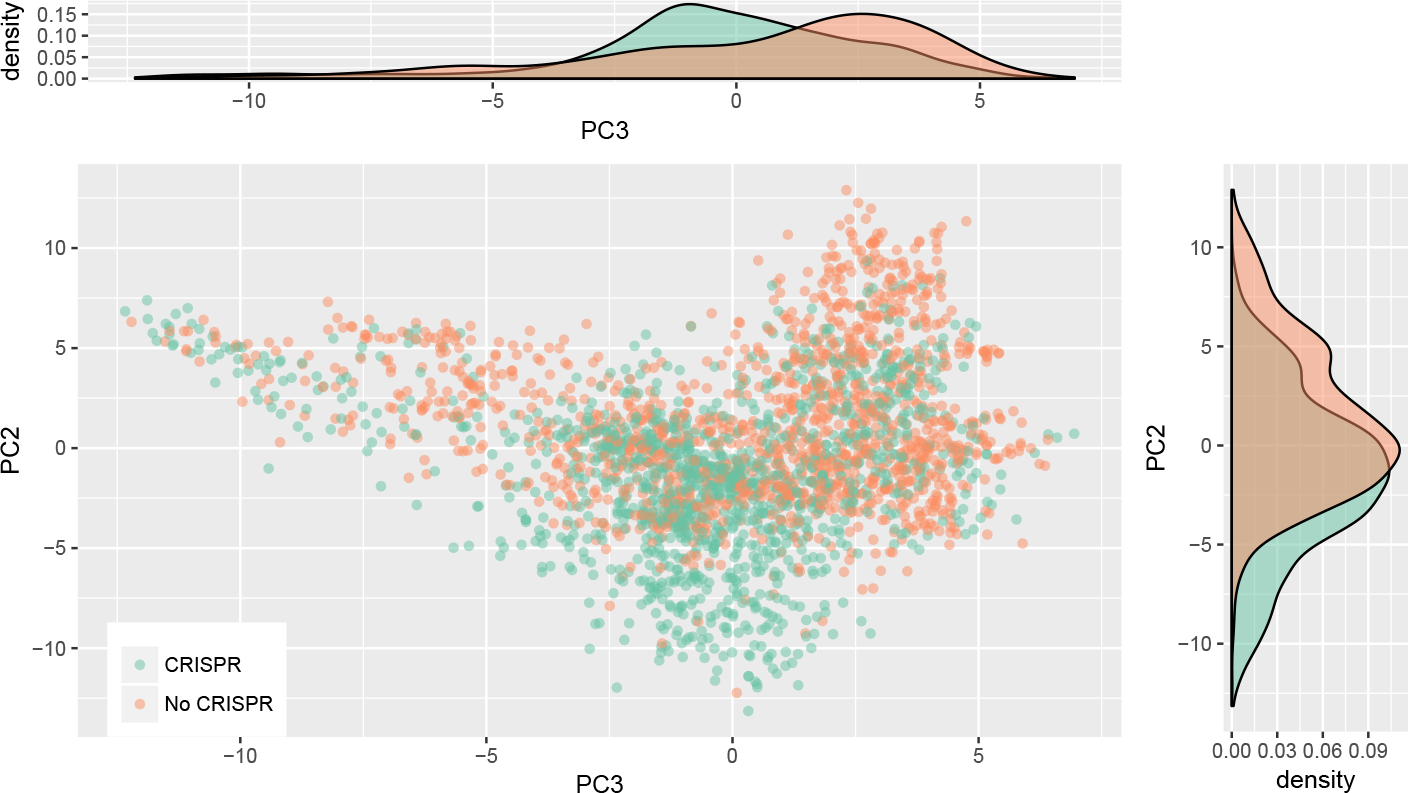
Organisms with CRISPR separate from those without in trait space. The second and third components from a PCA of the microbial traits dataset are shown. CRISPR incidence is indicated by color (green with, orange without), but was not included when constructing the PCA. Notice the separation of organisms with and without CRISPR along both components. Marginal densities along each component are shown to facilitate interpretation. See S1 Fig for the first component.

Second, we visualized the trait data using *t*-distributed stochastic neighbor embedding (t-SNE), which is a nonlinear method that can often pick up on more subtle relationships in a dataset (Fig 2; [26]). This method reveals a clustering of CRISPR-encoding microbes in trait space, further emphasizing that microbial immune strategy is influenced by ecological conditions. Because the axes of t-SNE plots are not easily interpretable, we mapped the top weighted traits from the PCA above (Table 1) onto the t-SNE reduced data (S2 Fig). Surprisingly, the most clearly aligned trait with CRISPR-incidence is having an obligately anaerobic metabolism.

**Figure 2:**
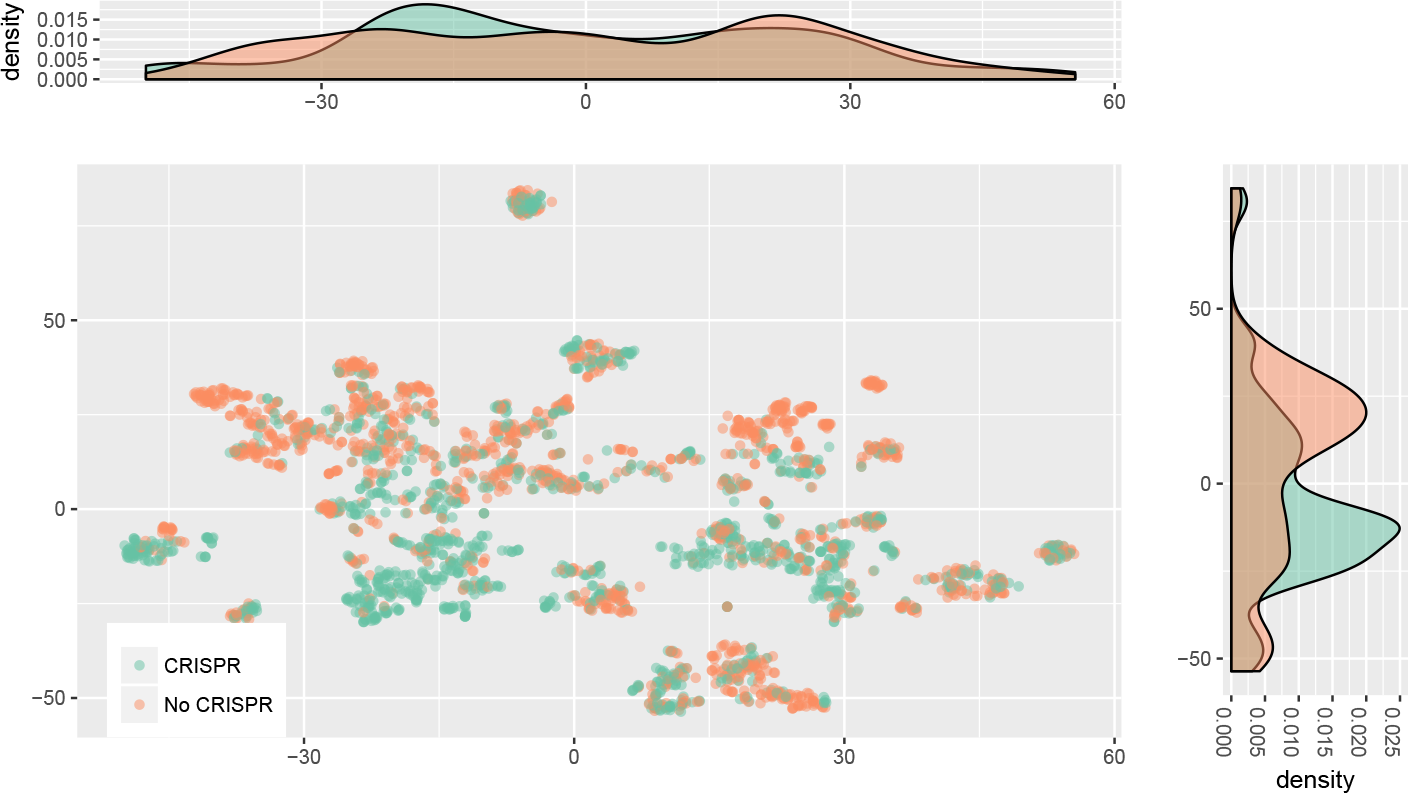
Organisms with CRISPR partially cluster in trait space away from those without. Two dimensional output of t-SNE dimension reduction on dataset. CRISPR incidence is indicated by color (green with, orange without), but was not included when performing dimension reduction. The axes of t-SNE plots have no clear interpretation due to the non-linearity of the transformation.

### Predicting CRISPR Incidence

The above unsupervised approaches (i.e. uninformed about the outcome variable, CRISPR) revealed patterns linking CRISPR incidence to microbial lifestyle. In order to further explore these patterns, and exploit them for their predictive ability, we applied several supervised prediction (i.e. trained with information about CRISPR incidence) methods to the data.

Unlike traditional statistical techniques focused on assigning *p*-values to particular input variables, with our machine learning approach we assessed model performance in terms of predictive ability. For unbiased error estimates, we chose an independent “test” set to withhold during the model fitting process and to be used only during model assessment. If we were able to effectively predict CRISPR incidence on this completely independent and previously unobserved dataset, then we concluded that our model encodes real information about which microbes possess CRISPR based on their traits. We then examined the structure of these models, and which variables play an outsize role in their performance, in order to select candidate traits associated with CRISPR incidence. Importantly, we chose the Proteobacteria as our test set because they represent a phylogenetically-independent group from our training set (see Methods).

All models we implemented showed improved predictive ability over a null model only accounting for the relative frequency of CRISPR among species (Cohen’s *κ* > 0; Table 2), indicating that there is some ecological signal in CRISPR incidence, though overall predictive performance was not overwhelming. Of these models the random forest (RF) model ranked highest, and did reasonably well (*κ* = 0.241). The percent incidences of CRISPR in the training (56%) and test sets (36%) are considerably different, which may have been difficult for these models to overcome. It is also possible that the Proteobacteria vary systematically from other phyla in terms of ecology and immune strategy, making them a particularly difficult (and thus conservative) test set. Nevertheless, the trait data clearly held some information about CRISPR incidence. We will primarily focus here on the RF model since it performed best, but see S1 Text for further discussion of the performance of our other models.

**Table 2:**
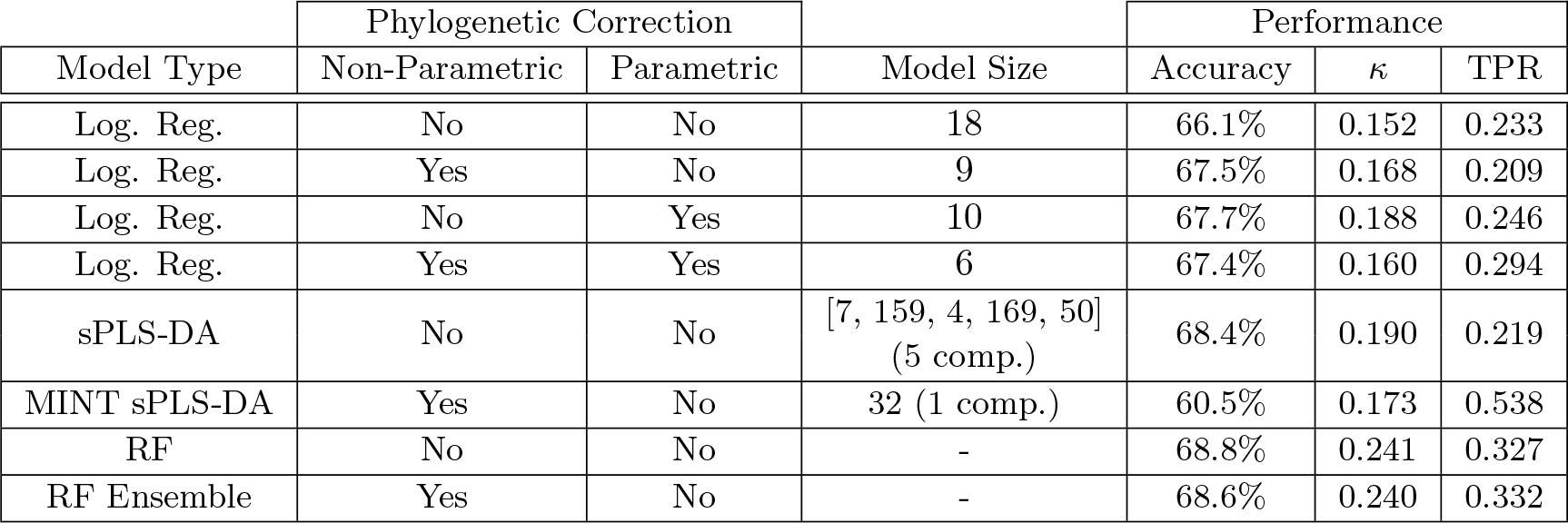
Predictive ability of models of CRISPR incidence on the Proteobacteria test set. Model size refers to number of variables chosen overall, or per-component in the case of the partial least squares models. Accuracy is measured as the total number of correct predictions over the total attempted and *κ* is Cohen’s *κ*, which corrects for uneven class counts that can inflate accuracy even if discriminative ability is low. Roughly, *κ* expresses how much better the model predicts the data than one that simply knows the frequency of different classes (*κ* = 0 being no better, *κ* > 0 indicating improved predictive ability). The true positive rate (TPR) is the number of correctly identified genomes having CRISPR divided by the total number of genomes having CRISPR in the test set. The non-parametric correction for phylogeny refers to our phylogenetically blocked folds, whereas the parametric correction refers to our use of phylogenetic logistic regression [61]. Observe that the RF model appears to perform best at prediction in general.

| Model Type | Phylogenetic Correction |  | Model Size | Performance |  |  |
| --- | --- | --- | --- | --- | --- | --- |
| | Non-Parametric | Parametric | | Accuracy | $\kappa$ | TPR |
| Log. Reg. | No | No | 18 | 66.1% | 0.152 | 0.233 |
| Log. Reg. | Yes | No | 9 | 67.5% | 0.168 | 0.209 |
| Log. Reg. | No | Yes | 10 | 67.7% | 0.188 | 0.246 |
| Log. Reg. | Yes | Yes | 6 | 67.4% | 0.160 | 0.294 |
| sPLS-DA | No | No | [7, 159, 4, 169, 50]<br>(5 comp.) | 68.4% | 0.190 | 0.219 |
| MINT sPLS-DA | Yes | No | 32 (1 comp.) | 60.5% | 0.173 | 0.538 |
| RF | No | No | - | 68.8% | 0.241 | 0.327 |
| RF Ensemble | Yes | No | - | 68.6% | 0.240 | 0.332 |

While each of our models revealed a distinct set of top predictors of CRISPR incidence, there was broad agreement overall (S1 Table, Fig 3, S3 Fig, and S4 Fig). Keywords indicating a thermophilic lifestyle (e.g. thermophilic, hot springs, hyperthermophilic, thermal springs) appeared across all models as either the most important or second most important predictor of CRISPR incidence. Keywords relating to oxygen requirement (e.g. anaerobic, aerobic) also appeared across nearly all models as top predictors, excluding only the two worst performing models (S1 Table). In the case of the RF and sPLS-DA models, oxygen requirement was always one of the top three predictors, and often the top predictor of CRISPR incidence (Fig 3, S3 Fig, S4 Fig, and S5 Fig). Other predictors that frequently appeared across model types included termite hosts (host insectstermites), the degradation of polycyclic aromatic hydrocarbons (PAH; metabolism pahdegrading), freshwater habitat (knownhabitats freshwater), and growth as filaments (shape filamentous). In general, the sPLS-DA, MINT sPLS-DA, RF, and RF ensemble models agreed with each other rather closely. Finally, we built an RF model using only traits related to temperature range, oxygen requirement, and thermophilic lifestyle (hot springs, thermal springs, hydrothermal vents). This temperature- and oxygen-only RF model outperformed all non-RF models (*κ* = 0.191). These traits alone appear to hold the majority of information about CRISPR incidence in the dataset.

**Figure 3:**
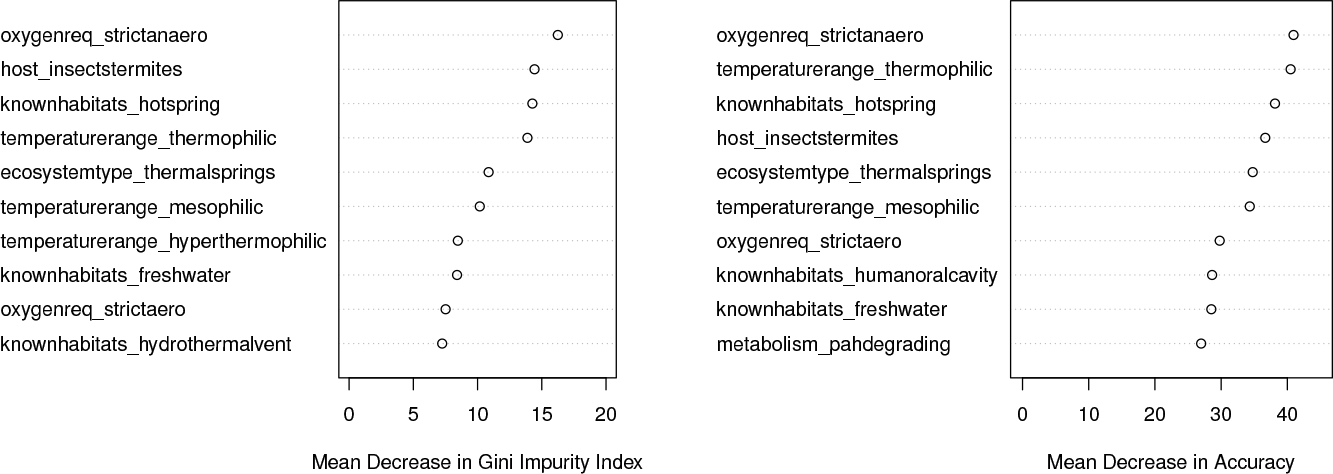
Importance of top ten predictors in the RF model. The mean decrease in accuracy measures the reduction in model accuracy when a variable is randomly permuted in the dataset. The Gini impurity index is a common score used to measure the performance of decision-tree based models (e.g. RF models). Briefly, when a decision tree is built the Gini impurity index measures how well separated the different classes of outcome variable are at the terminal nodes of the tree (i.e., how “pure” each of the nodes is). The mean decrease in Gini impurity measures the estimated reduction in impurity (increase in purity) when a given variable is added to the model. These importance scores are useful to rank variables as candidates for further study, but in themselves should not be taken as statistical support or effect sizes similar to those seen in linear regression. RF models may include non-linear combinations of variables, and therefore the contribution of any one variable is not as easily interpreted as with a linear model, a drawback of this approach. See S6 Fig for all predictor importances.

As an additional check that these candidate traits are associated with CRISPR incidence, we downloaded meta-data available from NCBI. We were able to reproduce the result that thermophiles strongly prefer CRISPR (92% with CRISPR as opposed to 49% in mesophiles, Fig 4a; [11, 12]). Though we have too few genomes categorized as psychrotolerant (35) or psychrophilic (14) to make any strong claims, these genomes seem to lack CRISPR most of the time, suggesting that CRISPR incidence decreases continuously as environmental temperatures decrease [10]. We were also able to confirm that, in agreement with our visualizations and predictive modeling, aerobes disfavor CRISPR immunity (34% with CRISPR) while anaerobes favor CRISPR immunity (67% with CRISPR, Fig 4b). This is true independent of growth temperature, with mesophiles showing a similarly strong oxygen-CRISPR link (S7 Fig). Overall, both oxygen (*χ*^2^ = 254.04, *p* < 2.2 × 10^−16^, categories with < 10 observations excluded) and temperature (*χ*^2^ = 98.86, *p* < 2.2 × 10^−16^, categories with < 10 observations excluded) had significant effects on incidence (for breakdown see Fig 4).

**Figure 4:**
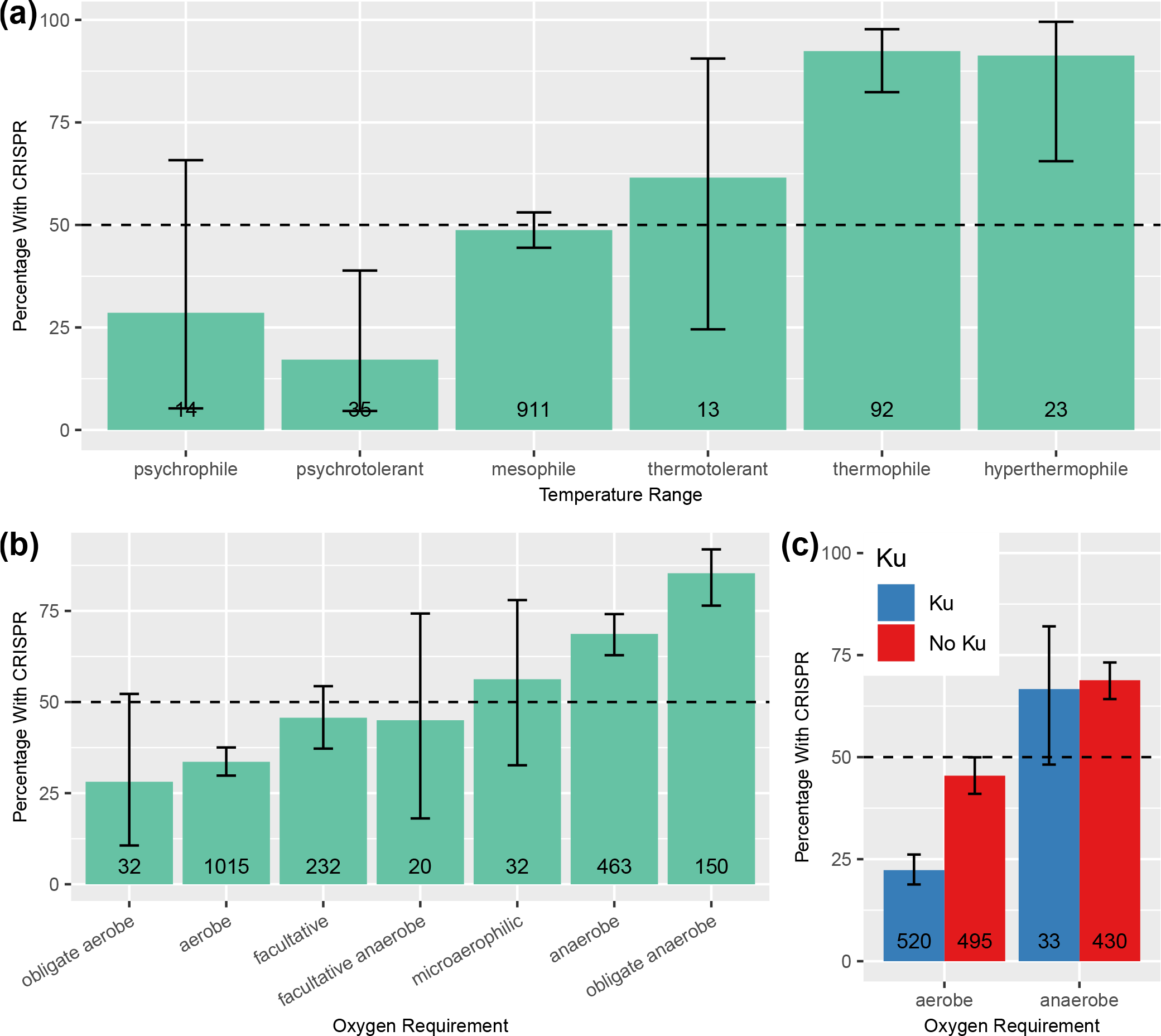
Temperature range and oxygen requirement are strong predictors of CRISPR incidence. Trait data taken from NCBI. (a) Thermophiles strongly favor CRISPR immunity, while mesophiles appear ambivalent. (b) Anaerobes favor CRISPR immunity, while aerobes tend to lack CRISPR and facultative species fall somewhere in between. (c) CRISPR and the Ku protein are negatively associated in aerobes but not anaerobes. Error bars are 99% binomial confidence intervals (non-overlapping intervals can be taken as evidence for a statistically significant difference at the *p* < 0.01 level). Total number of genomes in each trait category shown at the bottom of each bar. Categories represented 23 by fewer than 10 genomes were omitted.

Following previous suggestions that CRISPR incidence might be negatively associated with host population density and growth rate [11, 12, 15], and that this could be driving the link between CRISPR incidence and optimal temperature range, we sought to determine if growth rate was a major determinant of CRISPR incidence. The number of 16s rRNA genes in a genome is an oft used, if imperfect, proxy for microbial growth rates and an indicator of copiotrophic lifestyle in general [27, 28, 29]. While CRISPR-encoding genomes had slightly more 16s genes than CRISPR-lacking ones (3.1 and 2.9 on average, respectively), the 16s rRNA gene count in a genome was not a significant predictor of CRISPR incidence (logistic regression, *p* = 0.05248), although when correcting for phylogeny 16s gene count does seem to be significantly positively associated with CRISPR incidence (phylogenetic logistic regression, *m* = 0.06277, *p* = 6.651 × 10^−5^), the opposite of what we would expect if growth rate were driving the CRISPR-temperature relationship (though the effect is small).

As a secondary confirmation of the link between oxygen and CRISPR, we examined metagenomic data from the Tara Oceans Project [30], and found that across a large set of ocean metagenome samples CRISPR prevalence was inversely related to environmental oxygen concentration (S3 Text).

We also attempted to predict the number of CRISPR arrays in a genome given that that genome had at least one array, though this attempt was entirely unsuccessful (S4 Text).

### Predicting CRISPR Type

Each CRISPR system type is associated with a signature *cas* targeting gene unique to that type (*cas3*, *cas9*, and *cas10* for type I, II, and III systems respectively). There are many species in the dataset with *cas3* (605), but relatively few with *cas9* (160) and *cas10* (222), suggesting that the traits correlated with CRISPR incidence probably correspond primarily to type I systems (the dominance of type I systems has been noted previously [31]). We mapped the incidence of each of these genes onto the PCA we constructed earlier (see S1 Fig and Table 1), and found that *cas9* separates from *cas3* and *cas10* along the first component (Fig 5a). Broadly, this indicates that type II systems are more commonly found in host-associated than free-living microbes, the opposite of the other two system types.

**Figure 5:**
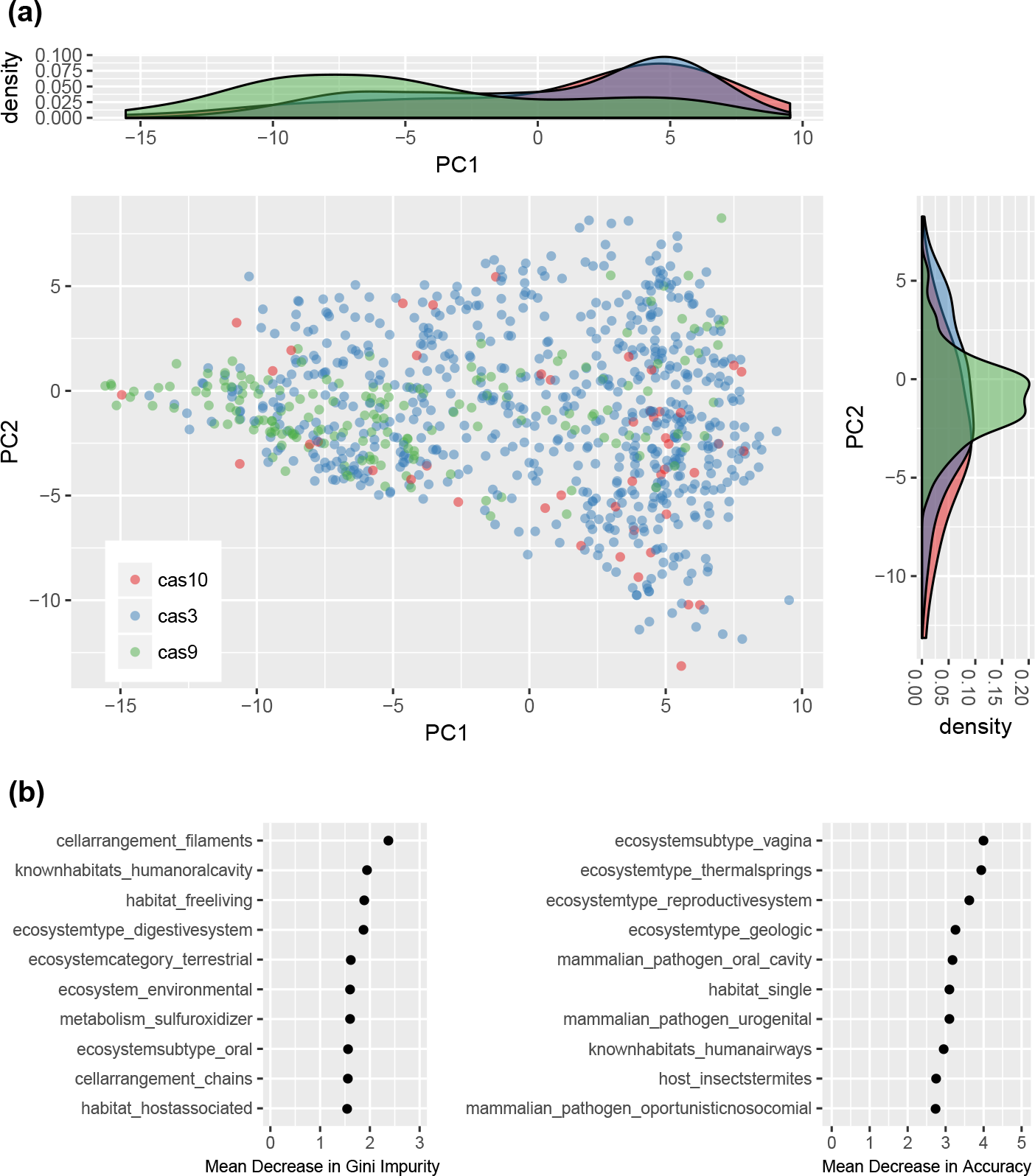
Type II CRISPR systems appear to be more prevalent in host-associated microbes. (a) The cas targeting genes associated with type I, type II, and type III systems (*cas3*, *cas9*, and *cas10* respectively) mapped onto the PCA in S1 Fig. Organisms without any targeting genes were omitted from the plot for readability. Recall from Table 1 that PC1 roughly corresponds to a spectrum running from host-associated to free-living microbes. (2) A variable importance plot from an RF model of *cas9* incidence. Observe that keywords related to a host-associated lifestyle appear many times.

We built an RF model of *cas9* incidence, with the Proteobacteria as the test set. Because our training set had so few cases of *cas9* incidence (10% of set), we performed stratified sampling during the RF construction process to ensure representative samples of organisms with and without *cas9*. Surprisingly, despite the extremely small number of organisms with *cas9* in the training and test sets (160 and 58 respectively), this model was accurately able to predict type II CRISPR incidence and had some discriminative ability (Accuracy = 93.0%, *κ* = 0.164), though it missed many of the positive cases (TPR = 0.172). This model also suggested that a host-associated lifestyle seems to be a major factor influencing the incidence of type II systems, with many of the top-ranking variables in terms of importance corresponding to keywords having to do with the split between host associated and free-living organisms (Fig 5b).

### NHEJ, CRISPR, and Oxygen

Recently, Bernheim et al. [21] demonstrated that the type II-A CRISPR system interferes with the NHEJ DNA repair pathway, leading to an inverse relationship between the presence of type II-A systems and the NHEJ pathway in microbial genomes. We hypothesized that this negative relationship between CRISPR and NHEJ might be more widespread across system types. We also hypothesized that this could explain the negative relationship between CRISPR and aerobicity we observe, since reactive oxygen species produced during aerobic respiration can induce double-strand breaks, thus selecting for the presence of NHEJ repair in aerobic organisms [32, 33]. We use the presence of Ku protein as a proxy for the NHEJ pathway, since this protein is central to the pathway.

There was a clear interaction between the presence of Ku and aerobicity on the incidence of CRISPR (Fig 4c). Using our full set of RefSeq genomes, we found a weak negative association between CRISPR and Ku incidence overall (Pearson’s correlation, *ρ* = −0.012; *χ*^2^ = 15.015, *p* = 1.067 × 10^−4^), but restricting only to aerobes the negative association between Ku and CRISPR was much stronger (Pearson’s correlation, *ρ* = −0.250, *p* = 9.109 × 10^−16^), whereas in anaerobes it was nonexistent (*ρ* = −0.023, *p* = 0.704). This pattern was consistent when correcting for phylogeny (S5 Text), and was true for both type I and III systems individually, though was not significant for type II systems of which there were fewer in the dataset S11 Fig.

Similar to our CRISPR analysis, we used PCA and an RF model to find if and where Ku-possessing organisms clustered in trait space. We found that the NHEJ pathway clusters strongly in trait space (S9 Fig), and is favored in soil-dwelling, spore-forming, aerobic microbes, consistent with expectations of where NHEJ will be most important [32, 33] (S10 Fig).

### Predicting RM Incidence

We attempted to determine whether RM systems were associated with similar microbial traits to those of CRISPR. In other words, are temperature and oxygen important determinants of whether a microbe has an intracellular immune system that degrades DNA in general, or are these drivers specific to CRISPR adaptive immunity? We built an RF model of restriction enzyme incidence using the same stratified sampling approach that we used for CRISPR system type. This model showed decent predictive ability (*κ* = 0.317). Overall, the correlation between variable importance scores for the CRISPR and restriction enzyme RF models was low (Pearson’s correlation, *ρ* = 0.169 for mean decrease in Gini Impurity Index, *ρ* = −0.0487 for mean decrease in accuracy, S13 Fig).

A mapping of restriction enzymes onto our trait PCA also contrasted with the mapping of CRISPR into trait-space (Fig S12 Fig). Because very few genomes lacked a restriction enzyme (97; in accordance with previous results [34]), we hesitate to make any strong claims, but the restriction enzyme-lacking organisms seem to be host associated (low values on PC1), thermophilic or anaerobic (low values on PC2), and solitary and motile (high values on PC3). We also found that the number of restriction enzymes was negatively associated with an anaerobic lifestyle (*m* = −4.53877, *p* = 2 × 10^−16^, phylogenetic linear regression), but not significantly associated with a thermophilic lifestyle after considering the effects of multiple testing (*m* = 1.51063, *p* = 0.03779, phylogenetic linear regression).

Despite this difference in trait associations, we confirmed that CRISPR incidence is positively associated with the number of restriction enzymes on a genome (6.23 with versus 4.36 without CRISPR, *t* = −9.038, *p* < 2.2× 10^−16^; *m* = 0.0676, *p* = 7.212 × 10^−13^, phylogenetic logistic regression; [34]) as well as whether or not a genome has any restriction enzymes (*χ*^2^ = 35.065, *p* = 3.189 × 10^−9^; *m* = 1.96127, *p* = 1.853 × 10^−14^, phylogenetic logistic regression) consistent with previous analyses [34].

## Discussion

We detected a clear association between microbial traits and the incidence of the CRISPR immune system across species. We found that two predictors were especially important for predicting CRISPR incidence, thermophilicity and aerobicity. The links between these two traits and CRISPR were confirmed with annotations from NCBI, and in the case of aerobicity with metagenomic data from the Tara Oceans Project (S3 Text; [30]). The relationship between temperature and CRISPR is well known [8, 9, 10], but we lend further support here by formally correcting for shared evolutionary history in our statistical analyses using both parametric and non-parametric approaches.

Previous theoretical models predict that CRISPR will be selected against in environments with dense and diverse viral communities [11, 12], since hosts are less likely to repeatedly encounter the same virus in such environments. These models in turn predict that in high-density host communities CRISPR will not be adaptive, since high host density leads to high viral diversity [11, 12], and that this might explain why potentially slow-growing thermophiles favor CRISPR immunity (as opposed to copiotrophic mesophiles). Our results show a marginal positive association between growth rate and CRISPR incidence, and that group-living microbes seem to favor CRISPR immunity, calling these prior viral diversity and density based explanations into question. Additionally, our analysis suggests that psychrophilic and psychrotolerant species disfavor CRISPR more strongly than mesophiles, which is not clearly explained or predicted by hypotheses based on host density.

We suspect that another factor could be affecting the degree of viral diversity that a host encounters, so that viral diversity is high in colder environments and low in hotter ones. Differences in dispersal limitation among viruses could lead to lower immigration rates in hot environments, as viral decay rates may be low at lower temperatures and high at higher temperatures [35], though this is highly speculative. We note that host dispersal rates are unlikely to affect the viral diversity seen by a host on average unless most of the host population is dispersing, an unrealistic expectation.

Surprisingly, we find that oxygen requirement appears to be just as important of a predictor of CRISPR incidence as temperature, and that this pattern is independent of any effect of temperature. Possibly, this association can be explained by inhibitory effects of CRISPR on NHEJ DNA repair. Type II-A CRISPR systems have been shown to directly interfere with the action of the NHEJ DNA repair pathway in prokaryotes [21]. Reactive oxygen species are produced during aerobic metabolism and can cause DNA damage [32], making NHEJ potentially particularly important in aerobes. Thus, if CRISPR interferes with the NHEJ repair pathway, and this pathway is important in aerobes, we would expect CRISPR incidence to be inversely related to the presence of oxygen. Our data showed a clear interaction between aerobicity and the NHEJ machinery in determining CRISPR incidence that suggests that the link between CRISPR and aerobicity may be mediated by the presence of the NHEJ pathway (Fig 4c). The Cas proteins share many structural similarities with proteins implicated in DNA repair, and in some cases prefer to associate with DSBs, and it is perhaps unsurprising that they appear to broadly inhibit the NHEJ pathway whose proteins may be competing for substrate [36].

As an alternative to our NHEJ hypothesis, could patterns in viral diversity explain the relationship between aerobicity and CRISPR incidence? The viral-decay hypothesis we proposed to explain the enrichment of thermophiles with CRISPR does not make sense in this context, since we might expect viruses to decay more readily in the presence of oxygen rather than under anoxic conditions. It is unclear to us why the viruses of anaerobes would be more dispersal limited. Nevertheless, if the viral communities infecting anaerobes were shown to be less diverse than those infecting aerobes this could also explain the increased incidence of CRISPR among these organisms.

We found no strong link between the incidence or number of RM systems on a genome and a thermophilic or anaerobic lifestyle, suggesting that the major drivers of CRISPR incidence are indeed CRISPR specific, consistent with our viraldiversity and NHEJ-inhibition hypotheses.

We were also able to show that CRISPR types vary in in terms of the environments they are found in, with type II systems appearing primarily in host-associated microbes. This phenomenon could be due in part to phylogenetic biases in the dataset, but our use of a phylogenetically independent test set lends credence to the overall trend. We have no clear mechanistic understanding of why *cas9* containing microbes tend to favor a host-associated lifestyle. Nevertheless this result may have practical implications for CRISPR genome editing, since it has recently been found that humans frequently have a preex-isting adaptive immune response to variants of the Cas9 protein [37]. We note that type I and III systems do not appear to have a strong link to host-associated lifestyles.

While our dataset spanned a broad phylogenetic range (with some notable exceptions such as the Candidate Phyla Radiation [38]), we had a limited number of microbial traits, which may have obscured some important CRISPR-trait associations. With the number of microbial genomes in public databases constantly expanding, so too should efforts to provide metadata about each of the organisms represented by those genomes. At least part of the problem lies in the lack of a universally accepted controlled vocabulary for microbial traits (similar to that provided by the Gene Ontology Consortium [39]), although some admirable attempts have been made [40, 41]. This would both facilitate the construction of more expansive trait databases, and would help deal with the issue of comparing traits that span many different scales.

The ecological drivers of microbial immune strategy are likely as diverse as the ever-increasing number of known prokaryotic defense systems [42, 43]. The database-centered approach we take here can be complemented by targeted studies examining shifts in immune strategy across environmental gradients (e.g., S3 Text) to provide a more fine-grained understanding of how microbial populations adapt to their local pathogenic and abiotic environments.

## Methods

### Data

For a schematic outlining the entire data compilation process see S14 Fig.

### Trait Data

We downloaded the ProTraits microbial traits database [44] which describes 424 traits in 3046 microbial species. These traits include metabolic phenotypes, preferred habitats, and specific behaviors like motility, among many others. ProTraits was built using a semi-supervised text-mining approach, drawing from several online databases and the literature. All traits are binary, with categorical traits split up into dummy variables (e.g. oxygen requirement listed as“aerobic”, “anaerobic”, and “facultative”). For each trait in each species, two “confidence scores” in the range [0, 1], are given, corresponding to the confidence of the text mining approach that a particular species does (*c*_+_) or does not (*c*_−_) have a particular trait.

We derived a single score (*p*) that captured the confidences both that a species does and does not have a particular trait. Assuming we want our score to lay in the interval [0, 1], such a score should be zero when we are completely confident that a species does not have a trait, one when we are completely confident that a species has a trait, and 0.5 when we are completely uncertain whether or not a species has a trait (i.e., equally confident that it does and does not have the trait). In the following formula, 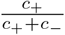 captures the relative confidence that a species does rather than does not have a trait, which we then scale by the overall maximal confidence (so that as overall confidence decreases the score shrinks towards 0.5)

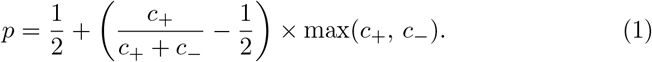

Many of the scores are missing for particular species-trait combinations (18%), indicating situations in which the text mining approach was unable to make a trait prediction. Our downstream analyses do not tolerate missing data, and so we imputed missing values using a random forest approach (R package missForest; [45]). There is a set of summary traits in the ProTraits dataset that were created de-novo using a machine learning approach, as well as a number of traits describing the growth substrates a particular species can use. We removed both summary and substrate traits from the dataset for increased interpretability (post-imputation; 174 traits remaining).

We note that the authors of ProTraits also used genomic data to help them infer strait scores, though we found that the exclusion of this data does not affect our overall outcome (S6 Text).

### Genomic Data and Immune Systems

For each species listed in the ProTraits dataset we downloaded a single genome from NCBI’s RefSeq database, with a preference for completely assembled reference or representative genomes. See S2 Text for a confirmation that our results are robust to the resampling of genomes. A number of species (333) had no genomes available in RefSeq, or only had genomes that had been suppressed since submission, and we discarded these species from the ProTraits dataset.

CRISPR incidence in each genome was determined using CRISPRDetect [46]. Additionally, data on the number of CRISPR arrays found among all available RefSeq genomes from a species were taken from Weissman et al. ([47]). We downloaded the REBASE Gold database of experimentally verified RM proteins and performed blastx searches of our genomes against this database [48, 49]. The distribution of E-values we observed was bimodal, providing a natural cutoff (*E* < 10^−19^).

To assess the ability of a microbe to perform non-homologous end-joining (NHEJ) DNA repair we used hmmsearch to search the HMM profile of the Ku protein implicated in NHEJ against all RefSeq genomes (E-value cutoff of 10^−2^/number of genomes; Pfam PF02735; [50, 51, 52]). We also used the annotated number of 16s rRNA genes in each downloaded RefSeq genome as a proxy for growth rate and the annotated *cas3*, *cas9*, and *cas10* genes as indicators of system type [53]. Where available as meta-data from NCBI, we also downloaded the oxygen (1949 records) and temperature requirements (1094 records) for the biosample record associated with each RefSeq genome.

### Phylogeny

We used PhyloSift to locate and align a large set of marker genes (738) found broadly across microbes, generally as a single copy [54, 55]. Of these marker genes, 67 were found in at least 500 of our genomes, and we limited our analysis to just this set. Additionally, eight genomes had few (< 20) representatives of any marker genes and were excluded from further analysis. We concatenated the alignments for these 67 marker genes and used FastTree (general-time reversible and CAT options; [56]) to build a phylogeny (S15 Fig).

### Visualizing CRISPR/RM Incidence

The size of the ProTraits dataset, both in terms of number of species and number of traits, and the probable complicated interactions between variables necessitate techniques that can handle complex, large scale data. To visualize the structure of microbial trait space and the distribution of immune strategies within that space we made use of two unsupervised machine learning techniques, principal component analysis (PCA) and *t*-distributed stochastic neighbor embedding (t-SNE, perplexity = 50, 5000 iterations; [26]).

PCA is a well-used technique in ecology that allows us to reduce the dimensionality of a dataset for effective visualization in two-dimensional space. Essentially, we collapse our trait dataset into two or three composite traits and observe whether species with a particular immune strategy tend to vary systematically in terms of where they fall in this “trait space”. A newer variant of this approach, t-SNE, performs a similar process, but unlike PCA allows for non-linear transformations of trait space. Therefore, local structure and non-linear interactions between traits in high dimensional space are preserved by t-SNE but often not captured by PCA [26]. On the other hand, t-SNE axes are less easily interpreted precisely because they represent non-linear rather than linear combinations of variables.

### CRISPR/RM Prediction from ProTraits

In order to predict the distribution of CRISPR and RM systems, we applied a number of supervised machine learning approaches to our dataset (see S16 Fig for a flow-chart describing the logic behind our model choices). In order to obtain accurate estimates of model performance, we initially set aside a portion of the data as a test set to be used exclusively in model assessment after all models were constructed (no fitting to this set). Because of the underlying evolutionary relationships in the data, we chose a test set that is phylogenetically independent of our training set. Alternatively, if we were to draw a test set at random from the microbial species we would risk underestimating our prediction errors due to non-independence of the training and test sets [57]. We chose the Proteobacteria as a test set because they are well-represented in the dataset (1139 species), ecologically diverse, and highly heterogeneous in terms of CRISPR incidence (S17 Fig). The remaining phyla were used to train our models.

First we built a series of linear models to classify species by immune strategy (CRISPR present or absent) using logistic regression. We had a large number of predictor variables (100+), which necessitated a model-selection approach in order to build a reasonably (and optimally) sized model. We used a forward selection algorithm to select the optimal set of predictors for each model size, with mean squared error under cross validation (CV) as our optimality criterion. We then selected model size by comparing BIC among these optimal models (i.e., selecting the model with the lowest score).

Similar to choosing a test set, care must be taken when performing CV on phylogenetically-structured data. CV assumes that when the data is partitioned into folds, each of these folds is independent of the others. If we draw species at random from a phylogeny, this assumption is violated, since the same hierarchical tree-structure will underlay each fold. Therefore, it is better to perform “blocked” CV than random CV [57], wherein folds are chosen based on divergent groups on the tree (e.g. phyla). If each group has diverged far enough in the past from the others, we can consider these folds to be essentially evolutionarily independent in terms of trait evolution (see S18 Fig for a conceptual example). Therefore blocked CV is essentially a non-parametric method (i.e., no explicit evolutionary model) to account for the non-independence arising from the shared evolutionary history between species. We use both random and blocked CV to build models. We clustered the data into blocked folds using the pairwise distances between tips on our tree (partitioning around mediods, pam() function in R package cluster; [58, 59]). A key assumption we make here is that our folds can be taken as independent from one another (i.e. no effect of shared evolutionary history). Since these clusters correspond roughly to Phylum-level splits, and since CRISPR and other prokaryotic immune systems are rapidly gained and lost over evolutionary time [60], we are comfortable making this assumption. We also repeated this analysis using phylogenetic logistic regression to more formally correct for phylogeny (R package phylolm; [61, 62]). Phylogenetic logistic regression is a more powerful method since it fits an explicit model of trait evolution, although it relies on the assumption that traits evolve according to the chosen model and can give misleading results otherwise.

Stepwise methods for variable selection, such as those used above (i.e., forward subset selection), are simple, computationally feasible, and easy to implement and interpret, but perform poorly when variables in the dataset covary with one another (i.e. multicollinearity; [63, 64]). As it so happens, the trait data used here exhibit strong multicollinearity (R package mctest; [65, 66]). Therefore, we sought out methods that deal well with this type of data, specifically partial least squares regression (PLS; [63]). Briefly, PLS combines features of PCA and linear regression to find the linear combination of predictors that maximizes the variance of the data in the space of outcome variables. We use a variant of PLS, sparse partial least squares discriminant analysis (sPLS-DA), where the “sparse” refers to a built-in variable selection process in the model-fitting algorithm and “discriminant analysis” refers to the fact that we are focused on a classification problem (i.e., presence vs. absence of a particular immune strategy; tune.splsda() and splsda() functions in R package mixOmics; [67, 68]).

We also attempt to ameliorate the effects of shared evolutionary history on our PLS model by using a philosophically similar approach to our blocked CV method above. Multivariate integrative (MINT) sPLS-DA is a variant of PLS that can account for systematic variation between groups of data when those groupings are known (e.g., our phylogenetically-blocked folds from above). It was originally developed for use in situations where multiple experiments testing the same hypothesis could show systematic biases from one another. In our case, the history of prokaryotic evolution is our experiment, and deep branching lineages are our replicates. We apply MINT sPLS-DA to the data, using the same blocked folds we used for CV (tune() and mint.splsda() functions in R package mixOmics; [68, 69]).

While regression provides easily interpretable trait weights and is computational efficient, in order to capture higher-order relationships between microbial traits we needed more powerful methods. Random forests (RF) are an attractive choice for our aims since they produce a readily-interpretable output and can incorporate nonlinear relationships between predictor variables [70]. We built an RF classifier on our training data from 5000 trees (otherwise default settings in R package randomForest; [71]). To prevent fitting to phylogeny, we took an ensemble approach which was similar in philosophy to our blocked CV and MINT sPLS-DA approaches above. Using the phylogenetically blocked folds defined above we fit five individual forests, each leaving out one of the five folds. We then weighted these forests by their relative predictive ability on the respective fold excluded during the fitting process (measured as Cohen’s *κ*; [72]). We predicted using our ensemble of forests by choosing the predicted outcome with the greatest total weight.

## Acknowledgments

JLW was supported by a GAANN Fellowship from the U.S. Department of Education and the University of Maryland. WFF was partially supported the U.S. Army Research Laboratory and the U.S. Army Research Office under Grant W911NF-14-1-0490. PLFJ was supported in part by NIH R00 GM104158.

## Conflict of Interest

The authors declare no conflict of interest.

